# The genomic and bioclimatic characterization of Ethiopian barley (*Hordeum vulgare* L.) unveils challenges and opportunities to adapt to a changing climate

**DOI:** 10.1101/2022.05.16.492093

**Authors:** Basazen F. Lakew, Leonardo Caproni, Seyoum A. Kassaw, Mara Miculan, Jemal Seid Ahmed, Simona Grazioli, Yosef Gebrehawaryat Kidane, Carlo Fadda, Mario Enrico Pè, Matteo Dell’Acqua

## Abstract

The climate crisis is impacting agroecosystems of the global South, threatening the food security of millions of smallholder farmers. Understanding the effect of current and future climates on crop agrobiodiversity may guide breeding efforts and adaptation strategies to sustain the livelihoods of farmers cropping in challenging conditions. Here, we combine a genomic and climatic characterization of a large collection of traditional barley varieties from Ethiopia, key to food security in local smallholder farming systems. We employ data-driven approaches to characterize their local adaptation to current and future climates and identify barley genomic regions with potential for breeding for local adaptation. We used a sequencing approach to genotype at high-density 436 barley varieties, finding that their genetic diversity can be traced back to geography and environmental diversity in Ethiopia. We integrate this information in a genome-wide association study targeting phenology traits measured in common garden experiments as well as climatic features at sampling points of traditional varieties, describing 106 genomic loci associated with local adaptation. We then employ a machine learning approach to link barley genomic diversity with climate variation, estimating barley genomic offset in future climate scenarios. Our data show that the genomic characterization of traditional agrobiodiversity coupled with climate modelling may contribute to the mitigation of the climate crisis effects on smallholder farming systems.

## Introduction

Since the 1960s, the green revolution heralded by Norman Borlaug’s work at CIMMITY enabled a marked increase in crop yields worldwide (Bailey-Serres, Parker, Ainsworth, Oldroyd, & Schroeder, 2019; Pingali, 2012). Thanks to a successful combination of genetic innovation, chemistry, and mechanization, the global cereal production has increased by 280 percent in the last 60 years (FAOSTAT, 2022) and supported the growth of the world population. The green revolution is also at the root the process of intensification of modern agriculture, where uncontrolled increased productivity per surface unit may lead to an increased use of inputs and therefore footprint of agricultural practices, a reduction of agrobiodiversity, and an impact on small-scale indigenous farming systems (Khoury et al., 2022). Indeed, conventional farming may provide higher yields but needs uniform management and external inputs including irrigation, fertilizers, and pesticides (Mondal et al., 2020). When optimal management conditions are not met, green revolution crop varieties can be outperformed by traditional varieties with local adaptation traits (Ricciardi, Mehrabi, Wittman, James, & Ramankutty, 2021). This is especially relevant in smallholder farming systems, which involve an estimated 570 million farms worldwide supporting the livelihoods of 2 billion people in highly diversified low-input cropping environments (Lowder, Skoet, & Raney, 2016). Here, low adoption rates are observed for modern improved varieties lacking adaptation to farming conditions as well as local culture (Acevedo et al., 2020), jeopardizing the impacts of breeders’ efforts in the face of yield advantages which are not realized in the use environments (Mancini et al., 2017).

The adaptive capacity of subsistence-based agricultural practices is depending on the severity and speed of climate change, but context matters as well. In challenging farming systems, the climate crisis exposes smallholder farmers to severe fluctuations that thereby magnify the challenges they must face to avoid losses (Cohn et al., 2017). In these contexts with high vulnerability, high economic damages resulting in food insecurity are very likely (Coronese, Lamperti, Keller, Chiaromonte, & Roventini, 2019). In the wake of climate change, new varieties that combine yield stability with local adaptation as well as appreciation by local users are paramount to achieve resilience and support food security (Simmonds, 1991). The Ethiopian cropping system is exemplative of this challenge. As a center of domestication and diversity of crops (Vavilov, 1951), Ethiopia still hosts highly diversified farming systems relying largely on smallholder farmers that maintain and cultivate thousands of landraces. This wealth of diversity is reflected by highly diversified landscapes, with 32 registered agroecological zones cultivated by approximately 80 million people (Bachewe & Taffesse, 2018). Previous studies on the Ethiopian landrace germplasm showed that the local agrobiodiversity is large but poorly represented in breeding, as in the case of durum wheat (Maccaferri et al., 2019) and barley (Milner et al., 2019). Barley (*Hordeum vulgare* L.) is the fifth most cultivated cereal in the country, grown by more than 4 million smallholder farmers on about one million hectares (Central Statistical Agency of Ethiopia, 2022; FAOSTAT, 2022). Ethiopian barley germplasm is markedly different from the international allele pool (Milner et al., 2019), and has potential for supporting breeding programs at local and international level (Jørgensen, 1992; Piffanelli et al., 2004), especially those targeting improved adaptation potential to enhance resilience to the climate crisis.

Until the recent past, barley has been cropped in Ethiopia in two seasons per year, the short rainy season *Belg* (February to April) and the main rainy season *Meher* (June to December). However, changing patterns of rainfall across the Horn of Africa makes it difficult to crop in two seasons (Wakjira et al., 2021). The Horn of Africa is among the most vulnerable regions in the continent where drastic changes in rain patterns are expected to have even more substantial impacts on food production in the future (Serdeczny et al., 2017). These projections are anticipated to coincide with low adaptive capacity as climate change intensifies stressors on small-scale agriculture (Morton, 2007). Today, breeding can operate in a big data dimension that connects genomics (Varshney et al., 2021), phenomics (Hickey, Chiurugwi, Mackay, & Powell, 2017), as well as climatic data and associated models describing current and projected climates (Yoder et al., 2014). When combining genomic information with remote sensing data and global descriptors of climatic conditions, it is possible to associate genetic diversity with environmental diversity at cultivation sites, identifying genomic loci associated with local adaptation towards specific climates (Capblancq, Fitzpatrick, Bay, Exposito-Alonso, & Keller, 2020). Genomic information can then be used to speed up the development of *climate-ready* varieties through the application of classical or innovative approaches, including genomic selection (Scheben, Yuan, & Edwards, 2016) and decentralized breeding (de Sousa et al., 2021).

In this study, we tap into the broad diversity of Ethiopian barley landraces to assess their genomic signatures for uniqueness and adaptation to current and future climates. We use a data-driven approach combining genomics, common garden experiments, and climate data analysis to describe the relation between barley diversity and the landscape. We use a machine learning approach to model barley genetic composition across climate and geography of Ethiopia; we use these approaches to look at projected climates and identify which area of current barley cultivation is more likely to suffer the effects climate change. We integrate this information in a genome-wide association study (GWAS) targeting phenology traits and current climate, identifying genomic loci associated to adaptation. We discuss how transdisciplinary approaches combining quantitative genetics and climate science may support barley improvement, unlocking the full potential of local landraces to foster a more sustainable and resilient agriculture.

## Materials and Methods

### Plant material and DNA extraction

Plant materials used in this study were derived from the Ethiopian Biodiversity Institute (EBI) barley landrace collection. Accessions were prioritized by completeness of passport data, and a core collection of 249 barley landraces was selected to cover the entire geographical and agroecological range of barley cultivation in Ethiopia. Landrace accessions were purified in an ear-to-row reproduction in the main season of 2016 at the experimental station of Holeta, Ethiopia (9.065 N, 38.459 E). When different morphological types were present in an EBI accession, this was split in as many accessions as types, resulting in 418 purified barley lines derived from landrace accessions. Lines derived from landraces were sided by 40 improved lines, representing all main barley improved lines released for cultivation in the country at the time of the assembly of the panel. Improved lines were sourced from National and Regional agricultural research centers. Full description of the plant materials used in this study is reported in Table S1. The full collection of 458 barley was germinated to extract genomic DNA at the molecular biology laboratory of the EBI, Addis Abeba, Ethiopia. DNA was extracted from 3-5 seedlings per line, pooled in equal quantity, with the GenElute Plant Genomic DNA Miniprep Kit (Sigma-Aldrich, St Louis, MO) following the manufacturer’s protocol. DNA was shipped to Italy and evaluated for quality and quantified using a Microplate photometer (Thermo Scientific, Milford, MA) and agarose gel electrophoresis at the Scuola Superiore Sant’Anna molecular laboratories.

### Sequencing and bioinformatic analysis

Genotyping was performed at IGA Tech sequencing services (Udine, Italy) with a double digestion Restrictionsite Associated DNA sequencing (ddRAD-seq) approach (Peterson, Weber, Kay, Fisher, & Hoekstra, 2012). Genomic libraries were prepared using *SphI* and *EcoRI* and sequenced on an Illumina HiSeq2500 instrument (Illumina, San Diego, CA) with V4 chemistry in paired-end 125-bp mode. Raw reads were demultiplexed with Stacks (Catchen, Hohenlohe, Bassham, Amores, & Cresko, 2013) and quality was assessed with FastQC tool (v 0.11.5). Reads of each sample were mapped against the *Hordeum vulgare* reference genome (MorexV3, DOI: 10.5447/ipk/2021/3) with BWA (Burrows-Weeler-Aligner v.0.7.12) using the MEM algorithm (Li, 2013). The products of the alignment were subsequently indexed with PicardTools and SAMtools (Li et al., 2009). The HaplotypeCaller algorithm, implemented in GATK and run in per-sample mode (McKenna et al., 2010), was used for variant identification. Variant calling was completed with GenotypeVCFstool (Danecek et al., 2011) to derive single nucleotide polymorphisms (SNPs). The resulting raw SNPs were filtered using GATK for the following criteria: QUAL < 30; QD < 2.0; MQ < 40.0; AF < 0.01; DP < 580; *SNP clusters* (> 3 variants within 5 bp windows).

### Genomic diversity

When not stated otherwise, all data analysis was performed in R (R Core Team, 2018). SNPs with minor allele frequency (MAF) above 5% were used to calculate pairwise linkage disequilibrium (LD) with the *r^2^* metrics in *R/Ldheatmap* (Shin, Blay, Mcneney, & Graham, 2006). LD decay was estimated on SNPs having MAF> 0.05, as function of physical distance according to the Hill and Weir equation (Hill & Weir, 1988). A threshold of r^2^ = 0.3 was used to estimate average LD decay for each chromosome. Genetic structure and diversity of the barley collection were studied using a set of SNP markers pruned by LD at a threshold of *r^2^* = 0.5, using PLINK (Chang et al., 2015) *indep-pairwise* function with a 150-variant window moving in 5-variant steps. A complete linkage agglomerative clustering, based on pairwise identity-by-state (IBS) distance, was run using PLINK with defaults. A pairwise dissimilarity neighbor joining (NJ) analysis was run in Tassel (Bradbury et al., 2007) to describe the phylogenetic relationship among lines. Genetic groupings of samples were detected with Discriminant Analysis of Principal Components (DAPC) (Jombart et al., 2010) using *R/adegenet* (Jombart, 2008). The diversity among samples was further described using Principal Component Analysis (PCA) and Admixture (Alexander & Lange, 2011).

### Spatial and climatic characterization

GPS coordinates of landrace sampling locations, where available, were projected onto the map of Ethiopia using *R/raster* (Hijmans, 2015). Altitudes were derived at each site based on GPS coordinates, using the CGIAR SRTM database at 90 m resolution (Reuter, Nelson, & Jarvis, 2007). Agroecological zones (AEZ) used in this study are defined by the Ministry of Agriculture of Ethiopia (*Agro-Ecological zones of Ethiopia*, n.d.). The cultivation area of barley was defined by the union of all AEZ polygons in which at least two sampling sites of barley landraces were present. Significant associations between DAPC genetic clusters and administrative regions, subregions and AEZs were assessed using Pearson’s Chi-squared test of independence.

Historical climate data for the study area was derived from the fifth generation of European Centre for Medium-Range Weather Forecasts (ECMWF) atmospheric reanalysis version (ERA5) data, released under the Copernicus Climate Change Service (Hersbach et al., 2020). The dataset covers over 30 years (1981-2010) at a spatial resolution of 0.25°x 0.25°. Climatological biases in temperature and precipitation across the East African region are reduced in ERA5 compared to the previous version of ERA-interim (Gleixner et al., 2020). Future climate projections were obtained from daily climate data extracted from a subset of 38 climate models among the CMIP5 dataset at the horizon of 2050 and 2070 for two Representative Concentration Pathway (RCP): RCP4.5 and RCP8.5. We followed a recent study to select the representative CMIP5 GCM models, as the number of genuinely independent ones in the CMIP5 archive is significantly less than the number of those submitted (Sanderson, Knutti, & Caldwell, 2015). After reviewing previous works on model similarity (IPCC, 2019) and quality metrics compared to historical observational data, we selected four relatively independent models: CESM1-BGC, CMCC-CM, MIROC5, and MPI-ESM-MR, from which we prepared ensembles for future projected climate. Climate extremes and small-scale variability are essential drivers in many climate change impact studies. However, the spatial resolution achieved by the global models is still insufficient to identify the fine structure of rainfall intensity correctly. Therefore, we employed a stochastic downscaling and bias technique, the Rainfall Filtered Autoregressive Model (RainFARM), which has been extended for application to long climate simulations (D’Onofrio, Palazzi, Von Hardenberg, Provenzale, & Calmanti, 2014; Rebora, Ferraris, von Hardenberg, & Provenzale, 2006). To achieve optimal data spatial and temporal adjustment, we accessed data of 150 weather station of the Ethiopian National Meteorology Agency (NMA), covering the entire barley cropping range.

Both historical and projected climate data were then derived only within the main barley growing season of Ethiopia (June - December), the *Meher* (Gissila, Black, Grimes, & Slingo, 2004; Segele & Lamb, 2005). The resulting historical and future climate data were used to derive 19 bioclimatic variables using *R/dismo* (Hijmans, Phillips, Leathwick, & Elith, 2011). Bioclimatic variables are biological meaningful indicators, often used in species distribution modelling; they represent seasonal trends and define seasonality and limiting climatic factors. Eleven variables (from bio 1 to bio11) are derived from temperature data while the remaining eight are related to precipitation (from bio12 to bio19). Collinearity among bioclimatic variables was then assessed with the *ensemble.VIF()* function in R/*BiodiversityR* (Kindt & Coe, 2005). Only variables with a Variance Inflation Factor (VIF) below 10 were retained for further analyses.

### Common garden-experiments and phenology data analysis

The barley collection was characterized in common garden experiments at two locations in Ethiopia, Arsi Negele (7° 21’ N, 38°242’ E) and Holeta (9° 00’ N, 38°30 E), during the *Meher* (June - December) of 2017 and the *Meher* of 2018. Samples were arranged using alpha lattice design with two replications per site, in plots consisting of four rows of 2.5 m in length and spaced 20 cm apart. Plots were fertilized with DAP and Urea as per the recommended rate of applications for the two sites and agronomic management was uniformly applied to all the genotypes. Two phenological traits were collected on full plots: days to 50% heading (DH) and days to 90% maturity (DM). Grain filling period (GFP, days from heading to maturity) was derived from DM and DH. Best linear unbiased predictions (BLUP) of measured traits were computed with R/ASReml (Gilmour, Gogel, Cullis, Welham, & Thompson, 2014) using the general model in Eq. (1):

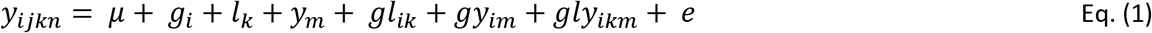

Where the observed phenotypic value is *y_ik_*, *μ* is the overall mean of the population, *g_i_* is the random effect for the *i^th^* genotype *g*, *l_k_* is the fixed effect for the *k^th^* location, *y_m_* is the random effect for the *m^th^* year. Interaction effects are considered for genotype, year, and location, and *e* is the error. For calculation of BLUPs within a single location and/or year, the data was sub-set and analyzed with a reduced model in accordance with Eq. (1). Broad-sense heritability (*H^2^*) was derived from the variance component estimates deriving from Eq. (1) as follows:

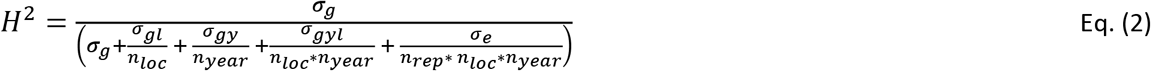

In Eq. (2), *σ_g_* is the variance component of genotypes,*σ_gl_* is the genotype by location variance, *σ_gy_* is the genotype by year variance, *σ_gly_* is the genotype by location by year variance, and *σ_e_* is the error variance. *n_loc_*, *n_year_*, *n_rep_* are the number of locations, years, and replications, respectively. For calculation of *H^2^* within locations and years, Eq. (2) was simplified accordingly.

### Estimation of Ethiopian barley genomic vulnerability to future climates

A Gradient Forest (GF) machine-learning regression tree-based algorithm was used to test which environmental variables best explained barley genetic variation across its cultivation area in Ethiopia. The GF, implemented in *R/gradientForest* (Ellis, Smith, & Pitcher, 2012), derives the importance, in terms of predictive power, of each bioclimatic variable in the change of allele frequencies to predict genetic composition across the climatic landscape. The GF was trained with environmental variables and with Moran’s eigenvector map (MEM) variables derived using the *dbmem*() function implemented in R/*adespatial* (S. Dray et al., 2012). MEMs are non-correlated eigenvectors of a spatial weighting matrix derived from geographic coordinates at sampling locations (Stéphane Dray, Legendre, & Peres-Neto, 2006; Griffith & Peres-Neto, 2006). A forest of 500 trees was built on each SNP as response variable, using climatic variables and MEMs as predictors. The GF model was then used to predict and measure the mismatch between the genetic composition under historical and future climate projections using the method of Fitzpatrick and Keller (2015). Genomic offset of Ethiopian barley landraces was calculated as the Euclidean distance between the allelic turnover under the historical and projected climate.

### Genome-wide associations

A GWAS was run on climatic variables and phenotypic BLUPs using a Fixed and random model Circulating Probability Unification (FarmCPU) (Liu, Huang, Fan, Buckler, & Zhang, 2016) implemented in R/rMVP (Yin et al., 2021). FarmCPU was run with correction for kinship, and the first 10 genetic PCs as covariates to consider population structure. Kinship was estimated using the VanRaden method (VanRaden, 2008). Both kinship and genetic PCs were calculated using the set LD-based pruned markers. The GWA was run on a set of high-quality SNP markers with MAF > 0.01. The thresholds to detect significant associations were set considering Bonferroni correction with alpha=0.05 and the whole SNP set and a False Discovery Rate (FDR) at 5% for multiple testing correction, computed with *R/qvalue* (Storey, Bass, Dabney, & Robinson, 2021).

The search for candidate genes was performed based on chromosome-specific LD as well as proximity to significant associations. Barley gene annotations were derived from the latest version of the barley reference genome (MorexV3, DOI: 10.5447/ipk/2021/3). Protein sequences of putative positional candidate genes were then used as queries against Araport11 reference proteome (Cheng et al., 2017).

## Results

### Genomic diversity in the collection

The sequencing of the collection yielded over 1.25 billion raw sequences. After variant calling and quality check, 436 barley lines and 47,492 genotyped SNPs were retained. LD in the collection decayed on average within 9.52 Mb, with variation across the seven chromosomes (Figure S1, table S2). A smaller set of 2,064 SNP markers was derived via LD-pruning to provide an unbiased representation of the diversity existing in the collection. Genetic diversity analyses depicted a complex genetic structure that could be best summarized by eleven genetic clusters (Figure S2, Figure S3), termed DAPC clusters 1-11. Improved barley varieties released for cultivation in Ethiopia were markedly different from most local landraces and consistently fell in DAPC cluster 2 and cluster 11 (Fig. 1). Landraces comprised all the other DAPC clusters and featured high genetic diversity; genetic structure was however low, as reported by the low variance explained by the PC axes (Fig. 1 a). Barley landraces having GPS coordinates at the sampling sites (n=383) span the entire geographical and AEZ range of barley cultivation in Ethiopia, from 1350 m a.s.l. to more than 3,780 m a.s.l. (Figure s4). The relation between the DAPC genetic clusters and the AEZs of origin of landraces showed that genetic groups are partially related to local cropping conditions. Ethiopian farmers cultivate barley in mid-highlands in sub-moist, moist, sub-humid and humid edaphic conditions, and DAPC clusters replaced one another across AEZs (Fig. 1 b). The DAPC cluster 8 and cluster 9 were mostly limited to sub-moist and sub-humid AEZs, respectively (Chi-square test p value < 0.05) (Figure S5). The distribution of the DAPC clusters could also be traced back to administrative borders of Ethiopia, which combine AEZs variation with historical and cultural diversity. DAPC cluster 8 was almost exclusively found in Tigray (Chi-square test p value < 0.05) (Fig. 1 c), and specifically *Misrakawi* (Eastern Tigray) and *Debubawi* (Southern Tigray) (Figure S5). DAPC cluster 10 and cluster 1 were mostly found in specific sub-regions of Amhara (*Debub Wello*) and Oromia (*Bale*), while DAPC cluster 9 and 3 were mainly found in *Semen Omo* and *Gurage* in the Southern Nations, Nationalities, and Peoples’ Region (SNNP). Cluster 3, in lesser extent, was also present in the sub-regions of *Hadiya* and *Keficho Shekicho* (Figure S6).

### Climatic and phenological diversity in the collection

The climate ensemble modeling revealed an increased trend for temperature-related bioclimatic variables across the Ethiopian barley cropping area at both scenarios and time horizons. Less variation was observed for precipitation related variables; however, some areas showed a projected increase in rainfall during the wettest months of cultivation of as much as 60% of the baseline (Figure S7). Temperature-related variables showed a decreasing signal in the southern and northeastern part of Ethiopia during the driest months of the barley growing season, especially under the most extreme RCP scenario (Figure S8).

To study the climatic diversity of the Ethiopian barley, a VIF approach was used to reduce multicollinearity of bioclimatic variables, identifying a set of nine variables that were used to derive the climatic features. Among the temperature-derived variables, bio2 (temperature range), bio3 (isothermality), bio4 (temperature seasonality) and bio9 (mean temperature of the driest quarter) were retained. Among the variables related to precipitation, bio12 (total precipitation), bio14 (precipitation of driest month), bio15 (precipitation coefficient of variation), bio18 (precipitation of warmest quarter) and bio19 (precipitation of coldest quarter), all associated with seasonality or limiting precipitation factors, were maintained.

The nine retained bioclimatic variables showed broad variation across the sampling area. When summarized by a PCA, the first bioclimatic PC was positively correlated with temperature range (bio2) and seasonality (bio4) as well as with precipitation seasonality (bio15) (Fig. 2 a) and explained 38.6 % of the climatic variance of the sites of collection of barley landraces. PC2, positively associated with precipitation variables (bio12 and bio19), accounted for 21.1 % of the bioclimatic variance, while PC3 explained 14.3 % of the variance, being negatively correlated with the temperature of the driest quarter (bio9). Climatic PC variables could tell apart DAPC clusters whose distribution pattern was associated with precipitation and temperature gradients (Fig. 2 b). DAPC cluster 9, mostly found in Tigray, is cultivated in warmer climates. DAPC cluster 5, not associated to any AEZ or administrative region, and cluster 1 appeared to be cultivated in wetter and drier conditions, respectively, when compared to the others. Some of the DAPC clusters, including cluster 2, seem to be cultivated in a broad combination of climatic conditions (Fig. 2 b).

The phenological characterization of the collection, carried out at two sites during the main season of 2017 and 2018, showed extensive variation in length of the plant cycle measured as days to heading (DH), days to maturity (DM), and grain filling period (GFP); BLUPs of these three traits are reported in Table S3). Landraces matured in a span of more than 30 days (mean=65.7, SD=7.5), but improved varieties required a longer maturing time (mean=72.8, SD=7.4). DH and DM had a strong genetic determination and were stable across locations and years, as reported by high broad sense heritability (*H^2^_B_*), while GFP to a lesser extent (Table S4). Phenology matched the genetic clustering observed in the collection. A PCA on phenology traits derived a PC1 highly correlated with DH and DM, explaining about two-thirds of the variance, while PC2 separated genotypes by GFP (Fig. 2 c). DAPC clusters are mostly ordered across PC1 and succeed one another in partially overlapping groupings. Several landraces are earlier than improved varieties, most notably cluster 1, 5,8, 9 and 10 which mature well before all the improved lines in the common garden experiment (Fig. 2 c).

### Outlook of barley cultivation in Ethiopia in a changed climate

The current extent and distribution of Ethiopian barley genomic diversity may not be suited to future climate scenarios. The GF analysis estimated the importance, in terms of predictive power, of the bioclimatic and spatial variables in relation to the genomic variation observed across the cropping area. Overall, the allelic turnover was best predicted by a bioclimatic indicator that describes variation of precipitation patterns (*i.e*. bio19), followed by a number of MEM variables representing spatial structure in the collection, and bio 9 (mean temperature of driest quarter) and bio12 (precipitation throughout the growing season) (Fig. 3 a). Predictors were used to estimate climate-driven genomic variation across the landscape (Fig. 3 c and d). Among the 23,674 SNPs tested as response variables, about 67% (n=15,869) were predicted by the model with *R*^2^ > 0.

Based on the current landscape-genome relations modeled through GF, we predicted the genomic composition at future RCP scenarios using projections derived from ensemble modeling climate data. We estimated barley future genomic offset as the difference between observed and expected genetic variation for each of the tested scenarios (Fig. 3 d). At all RCP-horizon combinations we observe high offset in the belt across the border that separates the region Afar from Tigray and Amhara, as well as in the western part of SNNP (Figure S9). The western part of Amhara and the region of Benishangul-Gumuz show medium offset (Fig. 3 d).

### Genome-wide associations and genomic signatures of adaptation

A GWAS was used to capture the genetic mechanisms underlying phenology and adaptation of Ethiopian barley. Twenty-four unique quantitative trait nucleotides (QTNs) were identified for DH and DM (Fig. 4 a and b) (Table S5). QTNs explained up to the 11.7% and 3.9% of variance of DH and DM, respectively. Some of the associations targeted well known flowering time genes in barley, but several others were unreported in literature. Among the genes implicated in flowering time variation in barley, the allelic variation of *FRIGIDA* and *VRN-H1* underly differences in phenology of Ethiopian barley. The same mapping approach was applied to the bioclimatic variables derived from historical climate, identifying 83 unique QTNs, 17 of which were associated with the two most important environmental predictors according to the GF model (bio19 and bio9) (Fig. 4 c and d) (Table S5). We found that a QTN was shared among DH and bio9 (SNP chr4H-88623874), tagging *FRIGIDA* (Fig. 4 a and d).

## Discussion

Our data shows that Ethiopian barley landraces are genetically different from most improved lines released for cultivation in the country (Fig. 1 a). This means that their allele pool is poorly explored by modern breeding, a finding in accordance with previous literature tapping into the world diversity of barley (Milner et al., 2019). Indeed, local breeding made limited use of Ethiopian agrobiodiversity and national tend to rely on international breeding materials rather than on traditional germplasm (Mengistu, Kidane, Fadda, & Pè, 2016). Still, some of the improved varieties included in this study showed some degree of overlap with landraces, possibly due to hybridizations by local breeding efforts or by mix-up at the field level, a feature not uncommon in material sourced from genebank collections (Sansaloni et al., 2020). The low use of the local allele pool to breed improved barley lines may leave behind variation relevant for adaptation to current and future climates.

Although high levels of admixture could be detected, as it is the case of other cereal landraces collections from the horn of Africa (Westengen et al., 2014), we reported a compelling association between the genomic makeup of landraces and Ethiopian AEZ and administrative regions (Fig. 1 b and c). Results show that the DAPC cluster 8 was almost limited to Tigray and to tepid sub-humid mid-highlands (SH3, Figure S4). This can be explained by local adaptation processes as well as with historical factors. Ethiopian smallholder farming makes little use of inputs, and largely depend on landraces that show adaptation to local environmental conditions.

The uniqueness of germplasm from Tigray was also reported on other crops, including teff (Woldeyohannes et al., 2022) and wheat (Di Falco, Chavas, & Smale, 2006). Tigray is more limiting in terms of barley potential production when compared to other regions of Ethiopia, and this aspect might have contributed to local selection for unique barley lineages. Historical and socio-economic influences may also have played an import role in shaping the distribution of the diversity on the landscape. Barley cultivation in the northern part of Ethiopia goes back to the Aksumites, one of the earliest civilizations in Horn of Africa (Harrower et al., 2019). Recent archeological evidence testifies and even earlier use of this crop in the region which corresponds to modern Tigray (D’Andrea, Perry, Nixon-Darcus, Fahmy, & Attia, 2018). Modern Tigrayan communities use barley to prepare *tihlo*, a typical dish based on barley which is not found in other parts of the country, and that may contribute to the uniqueness of the barley from this region.

The diversity in maturing time variation that we found in Ethiopian barley also reflects the extraordinary range of landscapes, farmers management practices and cultures. Phenology traits are quantitatively inherited (Table S4), interact with environmental factors and are of pivotal importance to maximize yield potential in target fields; well synchronized flowering and maturity time are necessary for achieving good harvest, especially in environments exposed to harsh conditions like terminal drought and heat (Jung & Müller, 2009). Indeed, control of phenology through breeding can reduce yield risks, for example achieving harvest earlier in the season.

Our climate projection analysis suggests that rain seasonality, as well as temperature, may change differently across the Ethiopian landscape. A remarkable difference in total precipitation, expected to increase more than 60%, may realize in the southwestern part of the country, in particular in *Keficho Shekicho* (Figure S7). As for temperature related variables, mean temperature of the driest quarter (bio9) showed a decreasing signal in the southern a northeastern part of Ethiopia under RCP 8.5 at both horizons, while no change signal was observed under RCP 4.5 (Figure S8). The ensemble approach used to generate projected climate showed comparable trends when compared to the historical dataset, suggesting that the projected climate data used in this study captured the ongoing climate trends across the region (Sanderson et al., 2015).

The combination of genomics, climatic data and common garden experiments was aimed at exploring the current and future climatic adaptation of Ethiopian barley germplasm. GF predictive models can significantly improve the ability to detect areas that are likely to be vulnerable (Rhoné et al., 2020); this approach represents a step forward if compared to other methods based on species distribution modelling (de Sousa, van Zonneveld, Holmgren, Kindt, & Ordoñez, 2019). The GF identified areas in which the extant landrace diversity of barley may not be well adapted to future climate. This can be put in relation with seasonal shifts we expect during the *Meher*.Under RCP8.5, the highest vulnerability for barley was predicted in northwestern SNNP (Fig. 3 d), in the administrative zone of *Keficho Shekicho*, at the border with *Gambela*, already among the warmest areas of Ethiopia (Degife, Zabel, & Mauser, 2021) and already marginal for barley production. Rather, the offset region bordering Afar (Fig. 3 d) is found in an area with high production of barley, and in close vicinity of areas of unique barley agrobiodiversity including Tigray, home of the cluster 8 described by this study (Fig. 1 c). The western part of Amhara as well as the region of Benishangul-Gumuz show medium offset to future climate scenarios; however, in this case, our dataset may not be representative, as the samples used in the analysis were collected in neighboring zones yet with same agroecological features. These results show that climate change will impact significant portions of the barley cropping areas in Ethiopia, especially those showing already hot and dry features that are projected to worsen. These findings suggest that breeding should focus on those areas exploiting local agrobiodiversity to enhance local adaptation of barley.

### Candidate genes and molecular targets for breeding

Modern breeding approaches could counteract the predicted vulnerability tapping into traits for local adaptation that are available in landraces. Our approach uncovered several QTNs associated with phenology and bioclimatic diversity, including the two bioclimatic variables having higher predictive power towards the GF. Some of the QTN tag genomic loci or genes already reported being associated with phenology and adaptation. A QTN at 527.9 Mb on Chr 5H, shared between DH and DM, is located 230 Kb downstream *VRN-H1* (Fu et al., 2005), homologous to *APETALA1/FRUITFUL (AP1/FUL*) of Arabidopsis. *VRN-H1* is a major player in flowering-time variation in barley, and its effect is conserved across different genetic backgrounds (Milner et al., 2019) and environmental conditions, with direct implications on yield (Francia et al., 2011; Tondelli et al., 2014). In cereals, transcription of *VRN1* increases with exposure to low temperatures (Danyluk et al., 2003), so that *VRN1* may also be related with recruitment of the floral-promoting potential of *AP1/FUL* genes to provide a low-temperature-induced flowering switch.

The most significant QTN for DH was found on chr4H at 88.6 Mb (Fig. 4 a), also significantly associated with mean temperature of driest quarter (bio9) (Fig. 4 d). This signal is located 185 Kb upstream of the gene model *HORVU.MOREX.r3.4HG0348870*, homolog to *FRIGIDA*. This gene has been extensively studied in Arabidopsis because of its role in flowering time variation (Noh & Amasino, 2003; Zhang & Jiménez-Gómez, 2020), having direct implications in floral transition. Latitudinal variation in flowering times in Arabidopsis is modulated by *FRIGIDA* (Stinchcombe et al., 2004), suggesting its key role in adaptation. DH and DM also share a significant SNP on chr7H; the signal lies 270 Kb upstream *enolase (ENO) (HORVU.MOREX.r3.7HG0647190*) that encodes for a protein homologous to Arabidopsis’ ENO2 (AT2G36530), that acts as positive regulator of cold-responsive gene transcription (Lee et al., 2002). In a recent work, Arabidopsis loss of function mutants for *ENO2* showed early flowering phenotypes (Ma et al., 2021) that reinforce the relevance of this candidate gene.

On the same chromosome chr7H we observed an association for precipitation of the coldest quarter (bio19) at 401.4 Mb, about 600 Kb from the gene model *HORVU.MOREX.r3.7HG0701970.1*, encoding a CALCIUM/CALMODULIN-REGULATED RECEPTOR-LIKE KINASE that shows high similarity with Arabidopsis CRLK1, involved in cold sensing and tolerance (Yang, Chaudhuri, Yang, Du, & Poovaiah, 2010). The same bioclimatic variable is associated with another signal on chr6H at 529.6 Mb. In this genomic region, 37Kb downstream the SNP, lies a *3-ketoacyl-CoA synthase (HORVU.MOREX.r3.6HG0621140*). This gene has been reported as an evolutionary conserved mechanism in barley and Arabidopsis for plant surfaces sensing of distantly related powdery mildews and involved in related defense mechanism from these fungi. At 231.9 Mb on chr6H we observe another signal of bio19 at about 606 Kb downstream *HORVU.MOREX.r3.6HG0582540*, which codes for a putative disease resistance protein; notably, the same chromosomic region it is also tagged by the top significant SNP associated with the temperature-related variable bio9. These finding may underlie patterns of adaptation of Ethiopian barley landraces to pathogens, a feature well known and already exploited by breeding (Jørgensen, 1992; Piffanelli et al., 2004), this time possibly in association with different rainfall regimes.

The GF results reinforce these findings. In several cases, loci associated to phenology and adaptive potential were also those best predicted by the variation of the climatic indicators. The association of DM on chr 2 with DM at 506.2 Mb, is also characterized by high *R^2^* towards the GF model (> 99^th^ percentile), meaning that its predictability through bioclimatic variables and geographic features is highly consistent. This signal lies within the gene body of *HORVU.MOREX.r3.2HG0172560*, encoding for a major facilitator superfamily transporter protein highly similar to Arabidopsis’ STP12 and STP10. This gene family is known to play a role in senescence, suggesting a possible role in the dynamics of maturation in barley and more. In Arabidopsis, STPs are also implicated in response mechanisms linked to a range of different environmental stressors like osmoregulation, salt tolerance, dehydration response and, notably, low temperature response (Büttner, 2007).

### Conclusions

Bridging genetic, climatic and phenological diversity under a genome-wide association framework allows the identification of genomic loci with potential for future breeding efforts addressing local adaptation and climate shifts. More studies are needed to further the characterization of genomic loci relevant for adaptation and validation of candidate genes, yet our results show that relevant adaptation traits exist in Ethiopian landraces, and that these genetic materials should be rapidly exploited by breeding to meet the needs of a changing climate. Leveraging local genetic diversity, the result of local adaptation and farmers knowledge, it may be possible to derive valuable options to foster new breeding models able to address smallholder farming system’s needs as well as put in place effective tailored strategies to protect farmer livelihoods and mitigate the effects climate change.

## Acknowledgements

This work was financed by Doctoral School in Agrobiodiversity at Scuola Superiore Sant’Anna in Pisa and the Deutsche Gesellschaft für Internationale Zusammenarbeit (GIZ) project ‘Strengthening cultivar diversity of barley and durum wheat to manage climate related risks and foster productivity in marginal areas of Ethiopia’. We thank Lorenzo Sena for the discussion of data and results. We wish to acknowledge the Ethiopian Biodiversity Institute for providing the genetic materials used in this study. The authors declare no conflict of interest.

## Author contributions

BF conducted data collection and data analysis with LC and MD. BF, YGK, and SA conducted field planning and phenotyping. MM managed sequencing data and produced SNPs. JSA conducted climate analysis. SG analyzed climatic and genomic data. MD, MEP, YGK and CF supervised the project. BF, LC, and MD drafted the manuscript. LC produced figures. All authors read the final version and approved submission.

## Data availability

Barley accessions are available upon request from the Ethiopian Biodiversity Institute (EBI, http://www.ebi.gov.et/). Raw DNAsequencing reads are available on the Short Read Archive at NCBI (https://www.ncbi.nlm.nih.gov/sra/), BioProject ID PRJNA841803; NCBI BioSample accessions from SAMN28618042 to SAMN28618521.

## References

Acevedo, M., Pixley, K., Zinyengere, N., Meng, S., Tufan, H., Cichy, K.,… Porciello, J. (2020). A scoping review of adoption of climate-resilient crops by small-scale producers in low-and middle-income countries. Nature Plants, 6(10), 1231–1241. https://doi.org/10.1038/s41477-020-00783-z Agro-Ecological zones of Ethiopia. (n.d.).

Alexander, D. H., & Lange, K. (2011). Enhancements to the ADMIXTURE algorithm for individual ancestry estimation. BMC Bioinformatics, 12(1), 246.https://doi.org/10.1186/1471-2105-12-246

Bachewe, F. N., & Taffesse, A. S. (2018). Supply response of smallholder households in Ethiopia. In The economics of teff: Exploring Ethiopia’s biggest cash crop (pp. 181–204). Washington, DC: International Food Policy Research Institute (IFPRI). https://doi.org/10.2499/9780896292833_08

Bailey-Serres, J., Parker, J. E., Ainsworth, E. A., Oldroyd, G. E. D., & Schroeder, J. I. (2019). Genetic strategies for improving crop yields. Nature, 575(7781), 109–118. https://doi.org/10.1038/s41586-019-1679-0

Bradbury, P. J., Zhang, Z., Kroon, D. E., Casstevens, T. M., Ramdoss, Y., & Buckler, E. S. (2007). TASSEL: Software for association mapping of complex traits in diverse samples. Bioinformatics, 23(19), 2633–2635. https://doi.org/10.1093/bioinformatics/btm308

Büttner, M. (2007). The monosaccharide transporter(-like) gene family in Arabidopsis. FEBS Letters, 581(12), 2318–2324. https://doi.org/10.1016/j.febslet.2007.03.016

Capblancq, T., Fitzpatrick, M. C., Bay, R. A., Exposito-Alonso, M., & Keller, S. R. (2020). Genomic Prediction of (Mal)Adaptation Across Current and Future Climatic Landscapes. Annual Review of Ecology, Evolution, and Systematics, 51(1), 245–269. https://doi.org/10.1146/annurev-ecolsys-020720-042553

Catchen, J., Hohenlohe, P. A., Bassham, S., Amores, A., & Cresko, W. A. (2013). Stacks: an analysis tool set for population genomics. Molecular Ecology, 22(11), 3124–3140. https://doi.org/10.1111/mec.12354

Central Statistical Agency of Ethiopia. (2022). CountrySTAT Ethiopia. Retrieved February 5, 2022, from http://ethiopia.countrystat.org/search-and-visualize-data/en/

Chang, C. C., Chow, C. C., Tellier, L. C., Vattikuti, S., Purcell, S. M., & Lee, J. J. (2015). Second-generation PLINK: rising to the challenge of larger and richer datasets. GigaScience, 4(1), 7.https://doi.org/10.1186/s13742-015-0047-8

Cheng, C.-Y., Krishnakumar, V., Chan, A. P., Thibaud-Nissen, F., Schobel, S., & Town, C. D. (2017). Araport11: a complete reannotation of the {Arabidopsis} thaliana reference genome. The Plant Journal: For Cell and Molecular Biology, 89(4), 789–804. https://doi.org/10.1111/tpj.13415

Cohn, A. S., Newton, P., Gil, J. D. B., Kuhl, L., Samberg, L., Ricciardi, V.,… Northrop, S. (2017). Smallholder Agriculture and Climate Change. Https://Doi.Org/10.1146/Annurev-Environ-102016-060946, 42, 347–375.https://doi.org/10.1146/ANNUREV-ENVIRON-102016-060946

Coronese, M., Lamperti, F., Keller, K., Chiaromonte, F., & Roventini, A. (2019). Evidence for sharp increase in the economic damages of extreme natural disasters. Proceedings of the National Academy of Sciences of the United States of America, 116(43), 21450–21455. https://doi.org/10.1073/PNAS.1907826116/SUPPL_FILE/PNAS.1907826116.SAPP.PDF

D’Andrea, A. C., Perry, L., Nixon-Darcus, L., Fahmy, A. G., & Attia, E. A. E. (2018). A Pre-Aksumite Culinary Practice at the Mezber Site, Northern Ethiopia. In Plants and People in the African Past (pp. 453–478). Cham: Springer International Publishing. https://doi.org/10.1007/978-3-319-89839-1_20

D’Onofrio, D., Palazzi, E., Von Hardenberg, J., Provenzale, A., & Calmanti, S. (2014). Stochastic rainfall downscaling of climate models. Journal of Hydrometeorology, 15(2), 830–843. https://doi.org/10.1175/JHM-D-13-096.1

Danecek, P., Auton, A., Abecasis, G., Albers, C. A., Banks, E., DePristo, M. A.,… Durbin, R. (2011). The variant call format and {VCFtools}. Bioinformatics, 27(15), 2156–2158. https://doi.org/10.1093/bioinformatics/btr330

Danyluk, J., Kane, N. A., Breton, G., Limin, A. E., Fowler, D. B., & Sarhan, F. (2003). TaVRT-1, a Putative Transcription Factor Associated with Vegetative to Reproductive Transition in Cereals. Plant Physiology, 132(4), 1849–1860. https://doi.org/10.1104/pp.103.023523

de Sousa, K., van Etten, J., Poland, J., Fadda, C., Jannink, J.-L., Kidane, Y. G.,… Dell’Acqua, M. (2021). Data-driven decentralized breeding increases prediction accuracy in a challenging crop production environment. Communications Biology. https://doi.org/10.1038/s42003-021-02463-w

de Sousa, K., van Zonneveld, M., Holmgren, M., Kindt, R., & Ordoñez, J. C. (2019). The future of coffee and cocoa agroforestry in a warmer Mesoamerica. Scientific Reports, 9(1), 8828.https://doi.org/10.1038/s41598-019-45491-7

Degife, A. W., Zabel, F., & Mauser, W. (2021). Climate change impacts on potential maize yields in Gambella Region, Ethiopia. Regional Environmental Change, 21(2). https://doi.org/10.1007/s10113-021-01773-3

Di Falco, S., Chavas, J. P., & Smale, M. (2006). Farmer Management of Production Risk on Degraded Lands: The Role of Wheat Genetic Diversity in Tigray Region, Ethiopia. Washington, DC. Retrieved from https://core.ac.uk/download/pdf/6289022.pdf

Dray, S., Pélissier, R., Couteron, P., Fortin, M.-J., Legendre, P., Peres-Neto, P. R.,… Wagner, H. H. (2012). Community ecology in the age of multivariate multiscale spatial analysis. Ecological Monographs, 82(3), 257–275. https://doi.org/10.1890/11-1183.1

Dray, Stéphane, Legendre, P., & Peres-Neto, P. R. (2006). Spatial modelling: a comprehensive framework for principal coordinate analysis of neighbour matrices (PCNM). Ecological Modelling, 196(3–4), 483–493. https://doi.org/10.1016/J.ECOLMODEL.2006.02.015

Ellis, N., Smith, S. J., & Pitcher, C. R. (2012). Gradient forests: calculating importance gradients on physical predictors. Ecology, 93(1), 156–168. https://doi.org/10.1890/11-0252.1

FAOSTAT. (2022). FAOSTAT database collections. Food and Agriculture Organization of the United Nations.

Fitzpatrick, M. C., & Keller, S. R. (2015). Ecological genomics meets community-level modelling of biodiversity: Mapping the genomic landscape of current and future environmental adaptation. Ecology Letters, 18(1), 1–16. https://doi.org/10.1111/ele.12376

Francia, E., Tondelli, A., Rizza, F., Badeck, F. W., Li Destri Nicosia, O., Akar, T.,… Pecchioni, N. (2011). Determinants of barley grain yield in a wide range of Mediterranean environments. Field Crops Research, 120(1), 169–178. https://doi.org/10.1016/j.fcr.2010.09.010

Fu, D., Szucs, P., Yan, L., Helguera, M., Skinner, J. S., Zitzewitz, J. von,… Dubcovsky, J. (2005). Large deletions within the first intron in VRN-1 are associated with spring growth habit in barley and wheat. Molecular Genetics and Genomics, (273), 54–65. https://doi.org/10.1007/s00438-004-1095-4

Gilmour, A. R., Gogel, B. J., Cullis, B. R., Welham, S. J., & Thompson, R. (2014). ASReml User Guide Release 4.1 Functional Specification. VSN International Ltd, Hemel Hempstead, HP1 1ES, UK Www.Vsni.Co.Uk.

Gissila, T., Black, E., Grimes, D. I. F., & Slingo, J. M. (2004). Seasonal forecasting of the Ethiopian summer rains. International Journal of Climatology, 24(11), 1345–1358. https://doi.org/10.1002/joc.1078

Griffith, D. A., & Peres-Neto, P. R. (2006). Spatial modeling in ecology: the flexibility of eigenfunction spatial analyses. Ecology, 87(10), 2603–2613. https://doi.org/10.1890/0012-9658(2006)87[2603:smietf]2.0.co;2

Harrower, M. J., Dumitru, I. A., Perlingieri, C., Nathan, S., Zerue, K., Lamont, J. L.,… Peterson, E. A. (2019). Beta Samati: discovery and excavation of an Aksumite town. Antiquity, 93(372), 1534–1552. https://doi.org/10.15184/aqy.2019.84

Hickey, J. M., Chiurugwi, T., Mackay, I., & Powell, W. (2017, September). Genomic prediction unifies animal and plant breeding programs to form platforms for biological discovery. Nature Genetics. Nature Publishing Group. https://doi.org/10.1038/ng.3920

Hijmans, R.., Phillips, S., Leathwick, J., & Elith, J. (2011). Package ‘dismo’. Retrieved from http://cran.r-project.org/web/packages/dismo/index.html

Hijmans, R. J. (2015). Raster: Geographic Data Analysis and Modeling. Retrieved from http://cran.r-project.org/package=raster

Hill, W. G., & Weir, B. S. (1988). Variances and covariances of squared linkage disequilibria in finite populations. Theoretical Population Biology, 33(1), 54–78. https://doi.org/10.1016/0040-5809(88)90004-4

IPCC. (2019). Climate Change and Land: an IPCC special report on climate change, desertification, land degradation, sustainable land management, food security, and greenhouse gas fluxes in terrestrial ecosystems.

Jombart, T. (2008). adegenet: a R package for the multivariate analysis of genetic markers. Bioinformatics, 24(11), 1403–1405. https://doi.org/10.1093/bioinformatics/btn129

Jombart, T., Devillard, S., Balloux, F., Falush, D., Stephens, M., Pritchard, J.,… Nei, M. (2010). Discriminant analysis of principal components: a new method for the analysis of genetically structured populations. BMC Genetics, 11(1), 94.https://doi.org/10.1186/1471-2156-11-94

Jørgensen, I. H. (1992). Discovery, characterization and exploitation of Mlo powdery mildew resistance in barley. Euphytica, 63(1-2), 141–152. https://doi.org/10.1007/BF00023919

Jung, C., & Müller, A. E. (2009). Flowering time control and applications in plant breeding. Trends in Plant Science, 14(10), 563–573. https://doi.org/10.1016/j.tplants.2009.07.005

Khoury, C. K., Brush, S., Costich, D. E., Curry, H. A., Haan, S., Engels, J. M. M.,… Thormann, I. (2022). Crop genetic erosion: understanding and responding to loss of crop diversity. New Phytologist, 233(1), 84–118. https://doi.org/10.1111/nph.17733

Kindt, R., & Coe, R. (2005). Tree diversity analysis: a manual and software for common statistical methods for ecological and biodiversity studies. World Agrofirestry Centre.

Lee, H., Guo, Y., Ohta, M., Xiong, L., Stevenson, B., & Zhu, J.-K. (2002). LOS2, a genetic locus required for cold-responsive gene transcription encodes a bi-functional enolase. The EMBO Journal, 21(11), 2692–2702. https://doi.org/10.1093/emboj721.11.2692

Li, H. (2013). Aligning sequence reads, clone sequences and assembly contigs with BWA-MEM.

Li, H., Handsaker, B., Wysoker, A., Fennell, T., Ruan, J., Homer, N.,… 1000 Genome Project Data Processing Subgroup. (2009). The Sequence Alignment/Map format and SAMtools. Bioinformatics (Oxford, England), 25(16), 2078–2079. https://doi.org/10.1093/bioinformatics/btp352

Liu, X., Huang, M., Fan, B., Buckler, E. S., & Zhang, Z. (2016). Iterative Usage of Fixed and Random Effect Models for Powerful and Efficient Genome-Wide Association Studies. PLOS Genetics, 12(2), e1005767.https://doi.org/10.1371/journal.pgen.1005767

Lowder, S. K., Skoet, J., & Raney, T. (2016). The Number, Size, and Distribution of Farms, Smallholder Farms, and Family Farms Worldwide. World Development, 87, 16–29.https://doi.org/10.1016/J.WORLDDEV.2015.10.041

Ma, X., Wu, Y., Ming, H., Liu, H., Liu, Z., Li, H., & Zhang, G. (2021). AtENO2 functions in the development of male gametophytes in Arabidopsis thaliana. Journal of Plant Physiology, 263, 153417. https://doi.org/10.1016/J.JPLPH.2021.153417

Maccaferri, M., Harris, N. S., Twardziok, S. O., Pasam, R. K., Gundlach, H., Spannagl, M.,… Cattivelli, L. (2019). Durum wheat genome highlights past domestication signatures and future improvement targets. Nature Genetics, 51(5), 885–895. https://doi.org/10.1038/s41588-019-0381-3

Mancini, C., Kidane, Y. G., Mengistu, D. K., Pè, M. E., Fadda, C., Dell’Acqua, M.,… Abate, G. (2017). Joining smallholder farmers’ traditional knowledge with metric traits to select better varieties of Ethiopian wheat. Scientific Reports, 7(1). https://doi.org/10.1038/s41598-017-07628-4

McKenna, A., Hanna, M., Banks, E., Sivachenko, A., Cibulskis, K., Kernytsky, A.,… DePristo, M. A. (2010). The genome analysis toolkit: A MapReduce framework for analyzing next-generation DNA sequencing data. Genome Research, 20(9), 1297–1303. https://doi.org/10.1101/gr.107524.110

Mengistu, D. K., Kidane, Y. G., Fadda, C., & Pè, M. E. (2016). Genetic diversity in Ethiopian Durum Wheat (Triticum turgidum var durum) inferred from phenotypic variations, 1–11. https://doi.org/10.1017/S1479262116000393

Milner, S. G., Jost, M., Taketa, S., Mazón, E. R., Himmelbach, A., Oppermann, M.,… Stein, N. (2019). Genebank genomics highlights the diversity of a global barley collection. Nature Genetics, 51(2), 319–326. https://doi.org/10.1038/s41588-018-0266-x

Mondal, S., Dutta, S., Crespo-Herrera, L., Huerta-Espino, J., Braun, H. J., & Singh, R. P. (2020). Fifty years of semi-dwarf spring wheat breeding at CIMMYT: Grain yield progress in optimum, drought and heat stress environments. Field Crops Research, 250, 107757. https://doi.org/10.1016/J.FCR.2020.107757

Morton, J. F. (2007). The impact of climate change on smallholder and subsistence agriculture. Proceedings of the National Academy of Sciences of the United States of America, 104(50), 19680–19685. https://doi.org/10.1073/pnas.0701855104

Noh, Y.-S., & Amasino, R. M. (2003). PIE1, an ISWI Family Gene, Is Required for FLC Activation and Floral Repression in Arabidopsis. The Plant Cell, 15, 1671–1682.https://doi.org/10.1105/tpc.012161

Peterson, B. K., Weber, J. N., Kay, E. H., Fisher, H. S., & Hoekstra, H. E. (2012). Double Digest RADseq: An Inexpensive Method for De Novo SNP Discovery and Genotyping in Model and Non-Model Species. PLoS ONE, 7(5), e37135.https://doi.org/10.1371/journal.pone.0037135

Piffanelli, P., Ramsay, L., Waugh, R., Benabdelmouna, A., D’Hont, A., Hollricher, K.,… Panstruga, R. (2004). A barley cultivation-associated polymorphism conveys resistance to powdery mildew. Nature, 430(7002), 887–891. https://doi.org/10.1038/nature02781

Pingali, P. L. (2012). Green revolution: Impacts, limits, andthe path ahead. Proceedings of the National Academy of Sciences of the United States of America, 109(31), 12302–12308. https://doi.org/10.1073/pnas.0912953109

R Core Team. (2018). R: A language and environment for statistical computing.

Rebora, N., Ferraris, L., von Hardenberg, J., & Provenzale, A. (2006). RainFARM: Rainfall downscaling by a Filtered Autoregressive Model. Journal of Hydrometeorology, 7(4), 724–738. https://doi.org/10.1175/JHM517.1

Reuter, H. I., Nelson, A., & Jarvis, A. (2007). An evaluation of void-filling interpolation methods for SRTM data. International Journal of Geographical Information Science, 21(9), 983–1008. https://doi.org/10.1080/13658810601169899

Rhone, B., Defrance, D., Berthouly-Salazar, C., Mariac, C., Cubry, P., Couderc, M.,… Vigouroux, Y. (2020). Pearl millet genomic vulnerability to climate change in West Africa highlights the need for regional collaboration. Nature Communications, 11(1). https://doi.org/10.1038/s41467-020-19066-4

Ricciardi, V., Mehrabi, Z., Wittman, H., James, D., & Ramankutty, N. (2021). Higher yields and more biodiversity on smaller farms. Nature Sustainability, 4(7), 651–657. https://doi.org/10.1038/s41893-021-00699-2

Sanderson, B. M., Knutti, R., & Caldwell, P. (2015). A representative democracy to reduce interdependency in a multimodel ensemble. Journal of Climate, 28(13), 5171–5194. https://doi.org/10.1175/JCLI-D-14-00362.1

Sansaloni, C., Franco, J., Santos, B., Percival-Alwyn, L., Singh, S., Petroli, C.,… Pixley, K. (2020). Diversity analysis of 80,000 wheat accessions reveals consequences and opportunities of selection footprints. Nature Communications, 11(1), 4572.https://doi.org/10.1038/s41467-020-18404-w

Scheben, A., Yuan, Y., & Edwards, D. (2016). Advances in genomics for adapting crops to climate change. Current Plant Biology, 6, 2–10.https://doi.org/10.1016/J.CPB.2016.09.001

Segele, Z. T., & Lamb, P. J. (2005). Characterization and variability of Kiremt rainy season over Ethiopia. Meteorology and Atmospheric Physics, 89(1–4), 153–180. https://doi.org/10.1007/s00703-005-0127-x

Serdeczny, O., Adams, S., Baarsch, F., Coumou, D., Robinson, A., Hare, W.,… Reinhardt, J. (2017). Climate change impacts in Sub-Saharan Africa: from physical changes to their social repercussions. Regional Environmental Change, 17(6), 1585–1600. https://doi.org/10.1007/s10113-015-0910-2

Shin, J.-H., Blay, S., Mcneney, B., & Graham, J. (2006). LDheatmap: An R Function for Graphical Display of Pairwise Linkage Disequilibria between Single Nucleotide Polymorphisms. Journal of Statistical Software, 16. https://doi.org/10.18637/jss.v000.i00

Simmonds, N. W. (1991). Selection for local adaptation in a plant breeding programme. Theoretical and Applied Genetics, 82(3), 363–367. https://doi.org/10.1007/BF02190624

Stinchcombe, J. R., Weinig, C., Ungerer, M., Olsen, K. M., Mays, C., Halldorsdottir, S. S.,… Schmitt, J. (2004). A latitudinal cline in flowering time in Arabidopsis thaliana modulated by the flowering time gene FRIGIDA. Proceedings of the National Academy of Sciences of the United States of America, 101(13), 4712–4717. https://doi.org/10.1073/pnas.0306401101

Storey, J., Bass, A. J., Dabney, A., & Robinson, D. (2021). Q-value estimation for false discovery rate control. Retrieved from http://github.com/jdstorey/qvalue

Tondelli, A., Francia, E., Visioni, A., Comadran, J., Mastrangelo, A. M., Akar, T.,… Pecchioni, N. (2014). QTLs for barley yield adaptation to Mediterranean environments in the “Nure” × “Tremois” biparental population. Euphytica, 197(1), 73–86. https://doi.org/10.1007/s10681-013-1053-5

VanRaden, P. M. (2008). Efficient Methods to Compute Genomic Predictions. Journal of Dairy Science, 91(11), 4414–4423. https://doi.org/10.3168/JDS.2007-0980

Varshney, R. K., Bohra, A., Yu, J., Graner, A., Zhang, Q., & Sorrells, M. E. (2021). Designing Future Crops: Genomics-Assisted Breeding Comes of Age. Trends in Plant Science, 26(6), 631–649. https://doi.org/10.1016/j.tplants.2021.03.010

Vavilov, N. I. (1951). The Origin, Variation, Immunity and Breeding of Cultivated Plants translated from Russian by K.Starr Chester. Chronica Botanica.

Wakjira, M. T., Peleg, N., Anghileri, D., Molnar, D., Alamirew, T., Six, J., & Molnar, P. (2021). Rainfall seasonality and timing: implications for cereal crop production in Ethiopia. Agricultural and Forest Meteorology, 310, 108633. https://doi.org/10.1016/J.AGRFORMET.2021.108633

Westengen, O. T., Okongo, M. A., Onek, L., Berg, T., Upadhyaya, H., Birkeland, S.,… Brysting, A. K. (2014). Ethnolinguistic structuring of sorghum genetic diversity in Africa and the role of local seed systems. Proceedings of the National Academy of Sciences of the United States of America, 111(39), 14100–14105. https://doi.org/10.1073/pnas.1401646111

Woldeyohannes, A. B., Iohannes, S. D., Miculan, M., Caproni, L., Ahmed, J. S., de Sousa, K.,… Dell’Acqua, M. (2022). Data-driven, participatory characterization of farmer varieties discloses teff breeding potential under current and future climates. BioRxiv, 2021.08.27.457623. https://doi.org/10.1101/2021.08.27.457623

Yang, T., Chaudhuri, S., Yang, L., Du, L., & Poovaiah, B. W. (2010). A calcium/calmodulin-regulated member of the receptor-like kinase family confers cold tolerance in plants. Journal of Biological Chemistry, 285(10), 7119–7126. https://doi.org/10.1074/jbc.M109.035659

Yin, L., Zhang, H., Tang, Z., Xu, J., Yin, D., Zhang, Z.,… Liu, X. (2021). rMVP: A Memory-efficient, Visualization-enhanced, and Parallel-accelerated tool for Genome-Wide Association Study. Genomics, Proteomics & Bioinformatics. https://doi.org/10.1016/J.GPB.2020.10.007

Yoder, J. B., Stanton-Geddes, J., Zhou, P., Briskine, R., Young, N. D., & Tiffin, P. (2014). Genomic Signature of Adaptation to Climate in Medicago truncatula. Genetics, 196(4), 1263–1275. https://doi.org/10.1534/genetics.113.159319

Zhang, L., & Jiménez-Gómez, J. M. (2020). Functional analysis of FRIGIDA using naturally occurring variation in Arabidopsis thaliana. The Plant Journal, 103(1), 154–165. https://doi.org/10.1111/tpj.14716

