## Supplemental Materials for "The genomic and bioclimatic characterization of Ethiopian barley (*Hordeum vulgare* L.) unveils challenges and opportunities to adapt to a changing climate"

**This file includes:**

Supporting Figure S1 to S9

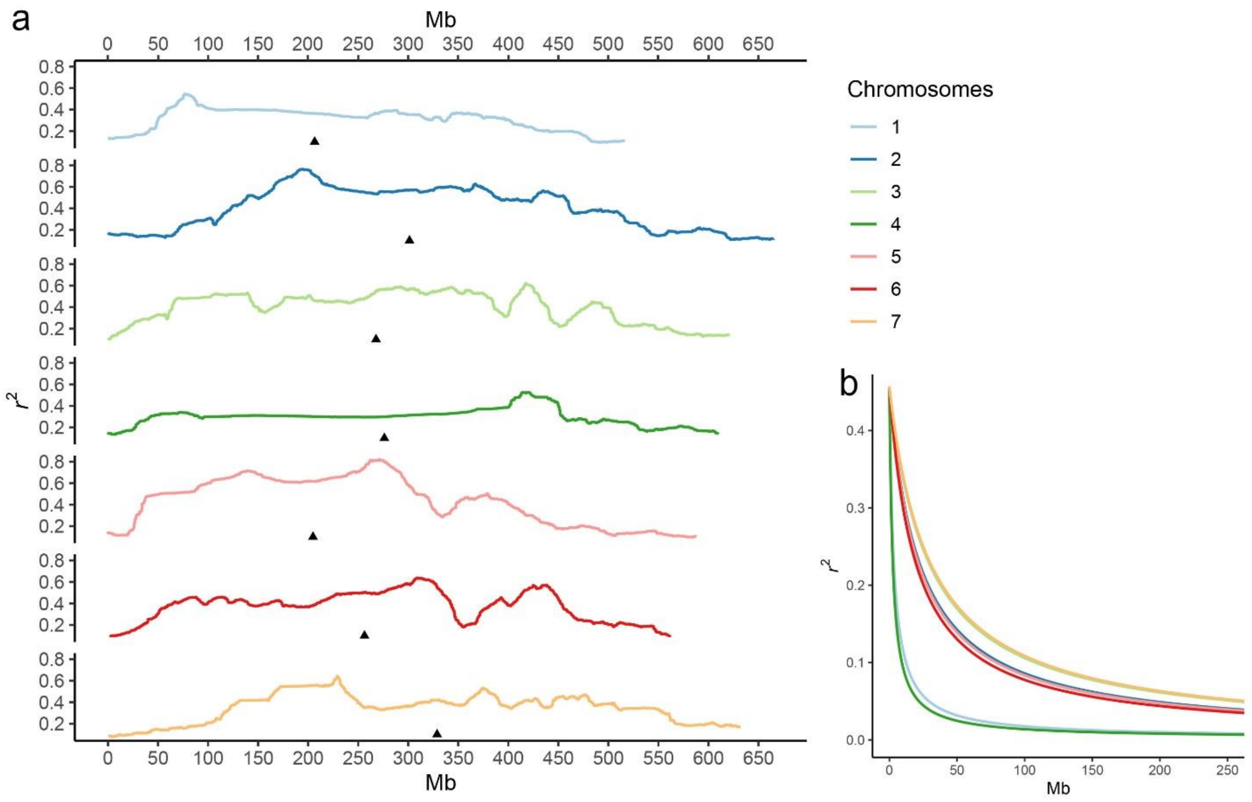


**Figure S1**. **a)** Genome-wide Linkage disequilibrium (LD) based on 436 barley genotypes sourced from EBI. The lines represent LD measures as a rolling window across the chromosome and are colored according to legend. Black arrowheads indicate the position of centromeres. **b)** LD decay as a function of physical distance of markers. Mb stand for million basepairs.

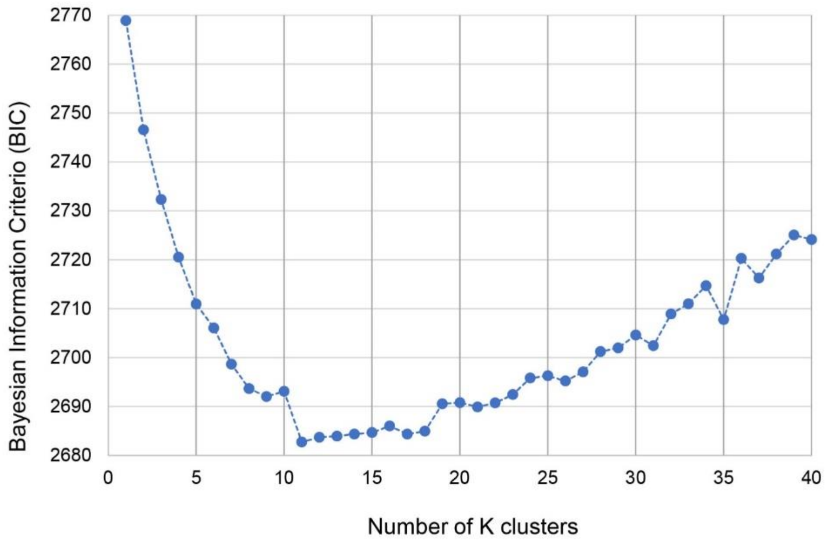


**Figure S2.** Predictive accuracy of the Discriminant Analysis of Principal Components (DAPC) procedure: Bayesian Information Criterion (BIC) values for different values of DAPC clusters.

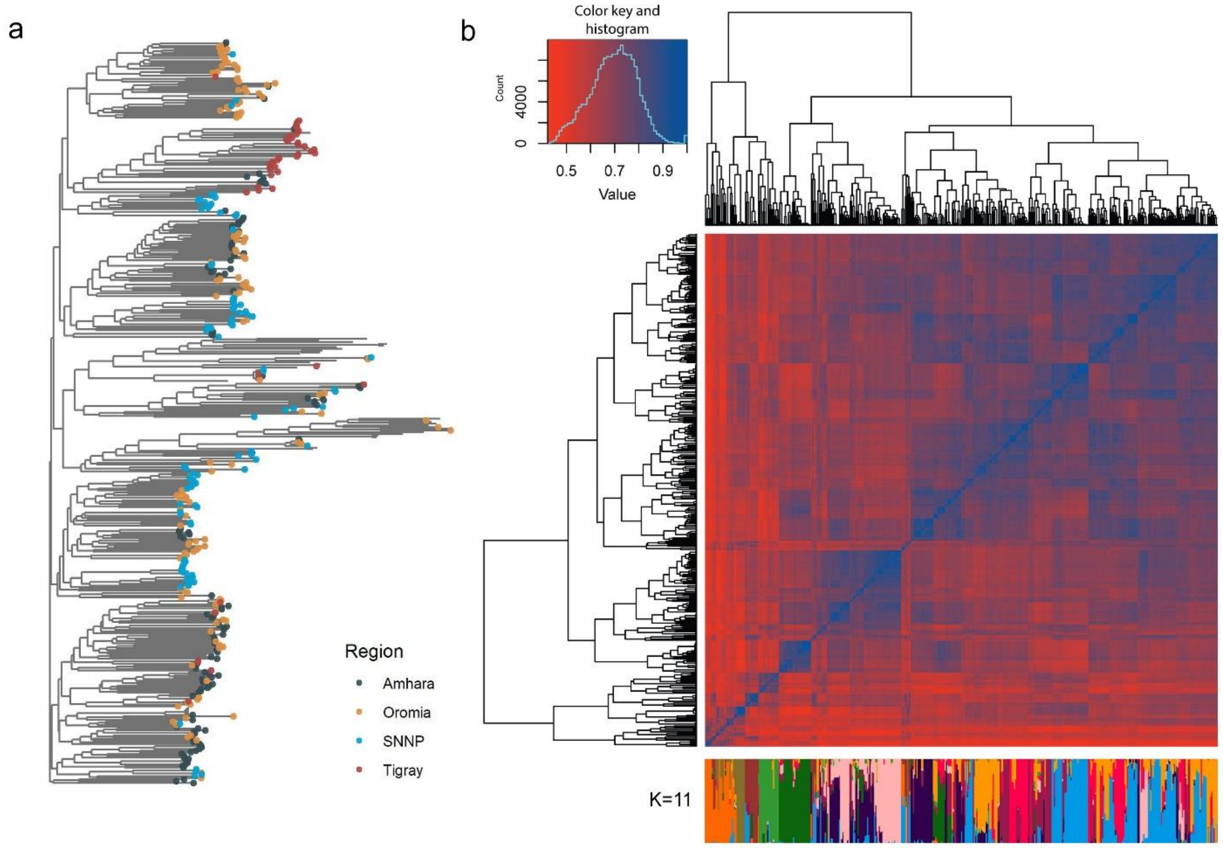


**Figure S3.** Genetic diversity analysis of 436 barley cultivated materials based on a set of 2,064 LD-pruned SNP markers: a) pairwise dissimilarity neighbor joining (NJ) tree, tips are colored by administrative region of origin of the landraces b) top: complete linkage agglomerative clustering, based on pairwise identity-by-state distance, bottom: ADMIXTURE result at K =11.

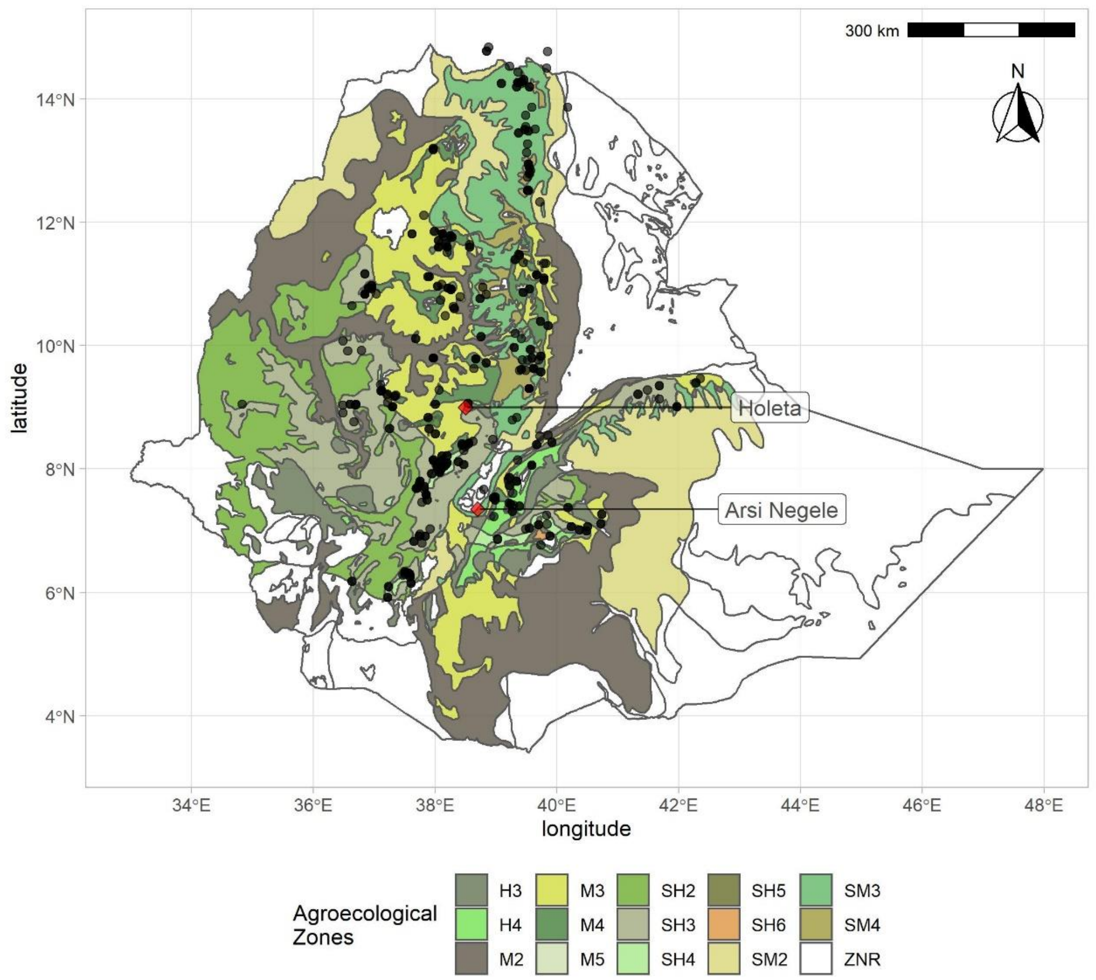


**Figure S4**. Sampling location of the barley accessions (N=249) across the agroecological zones of Ethiopia. H3, tepid humid mid-highlands; H4, cool humid mid-highlands; M2, warm moist lowlands; M3, tepid moist mid-highlands; M4, cool moist mid-highlands; SH2, warm sub-humid lowlands; SH3, tepid sub-humid mid-highlands; SH4, cool sub-humid mid-highlands; SH5, Cold sub-humid sub-afro-alpine to afro-alpine;SH6 Very cold sub-humid sub-afro to afro- alpine; SM2, warm sub-moist lowlands; SM3, tepid sub-moist mid-highlands; SM4, cool sub-moist mid-highlands; ZNR, zone not relevant (no hits). Common garden experiment locations are shown with red diamonds.

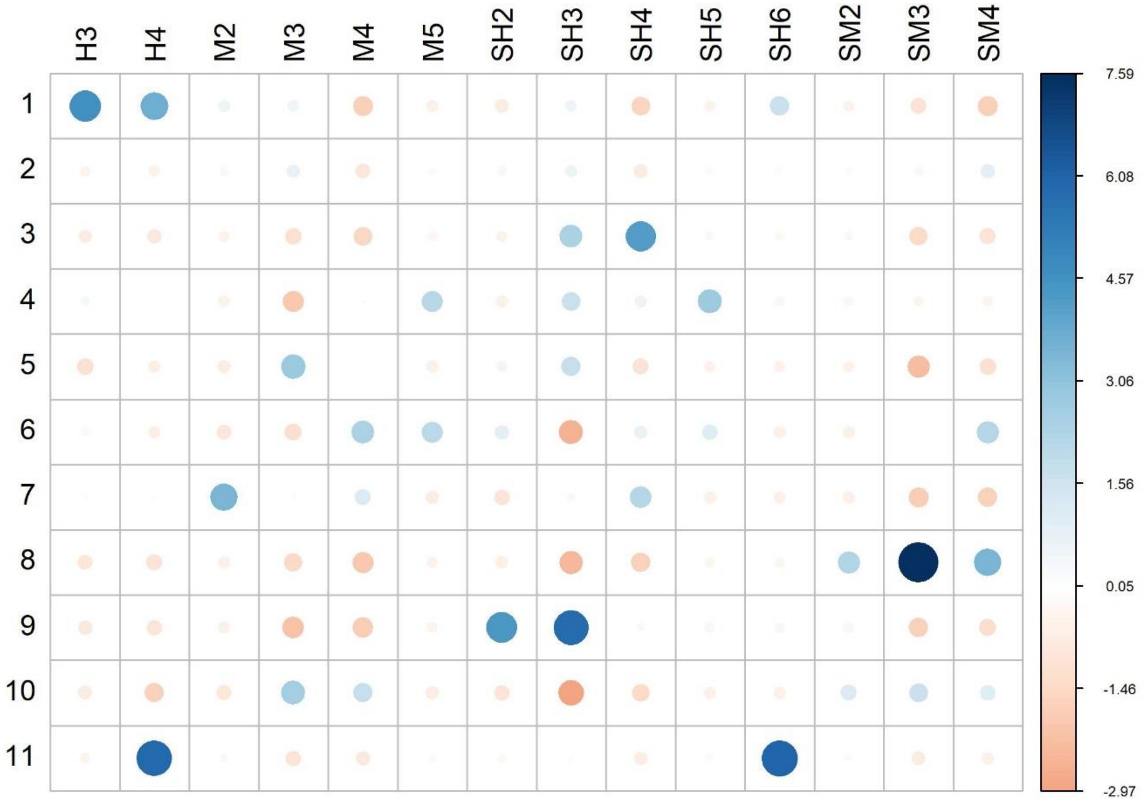


**Figure S5.** Residual plot for the Pearson's Chi-squared test of independence between genetic clusters and Agroecological zones of Ethiopia. H3, tepid humid mid-highlands; H4, cool humid mid-highlands; M2, warm moist lowlands; M3, tepid moist mid-highlands; M4, cool moist mid-highlands; SH2, warm sub-humid lowlands; SH3, tepid sub-humid mid-highlands; SH4, cool sub-humid mid-highlands; SH5, Cold sub-humid sub-afro-alpine to afro-alpine; SH6 Very cold sub-humid sub-afro to afro- alpine; SM2, warm sub-moist lowlands; SM3, tepid sub-moist mid-highlands; SM4, cool sub-moist mid-highlands.

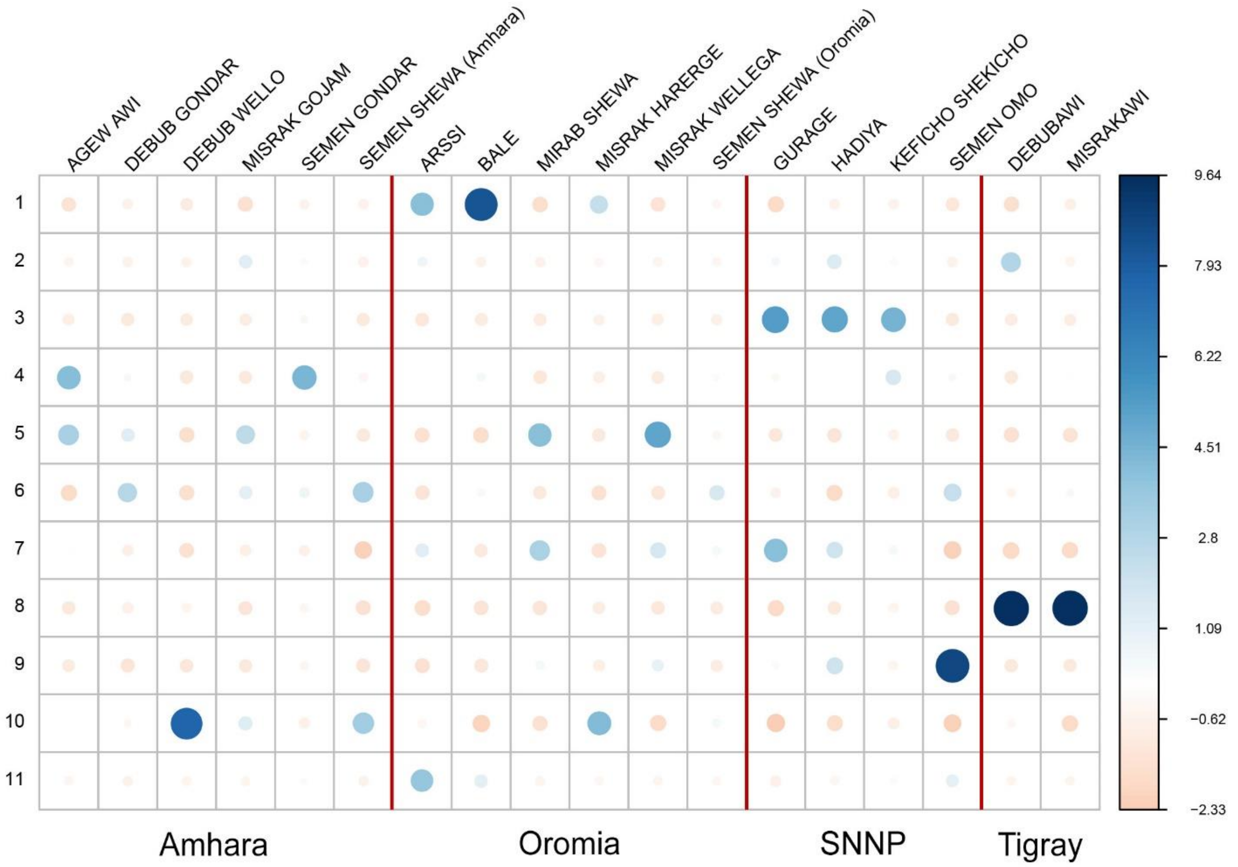


**Figure S6.** Residual plot for the Pearson's Chi-squared test of independence between genetic clusters and Ethiopian subregions (zones). Circle colors and sizes indicate the relative contribution of each cell to the Chi-square score. Dark blue and large circles indicate higher effect size.

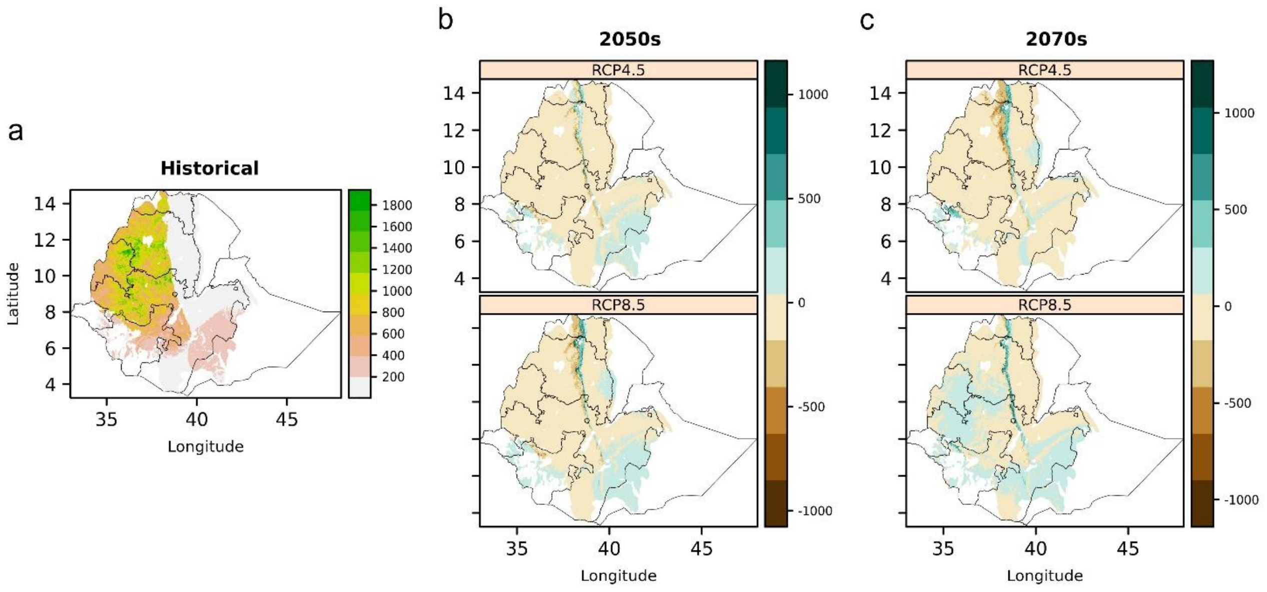


**Figure S7.** Precipitation of Coldest Quarter during the barley growing season in Ethiopia (bio19). a) Representation of historical data (mm during the growing season) in the period of 1981-2010. b) expected change at the horizon of 2050 at RCP 4.5 (top) and 8.5 (bottom) and c) at the horizon of 2070 at RCP 4.5 (top) and 8.5 (bottom), expressed in mm.

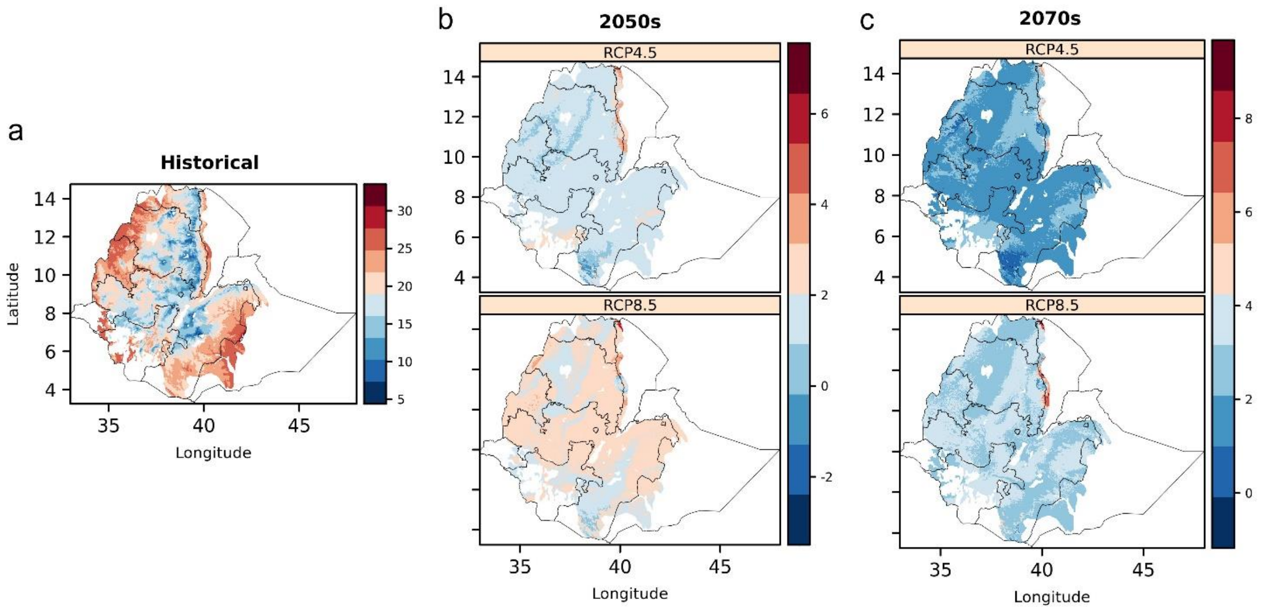


**Figure S8.** Mean Temperature of Driest Quarter (bio9). during the barley growing season in Ethiopia. a) Representation of historical data (°C) in the period of 1981-2010. b) expected change at the horizon of 2050 at RCP 4.5 (top) and 8.5 (bottom) and c) at the horizon of 2070 at RCP 4.5 (top) and 8.5 (bottom), expressed in °C.

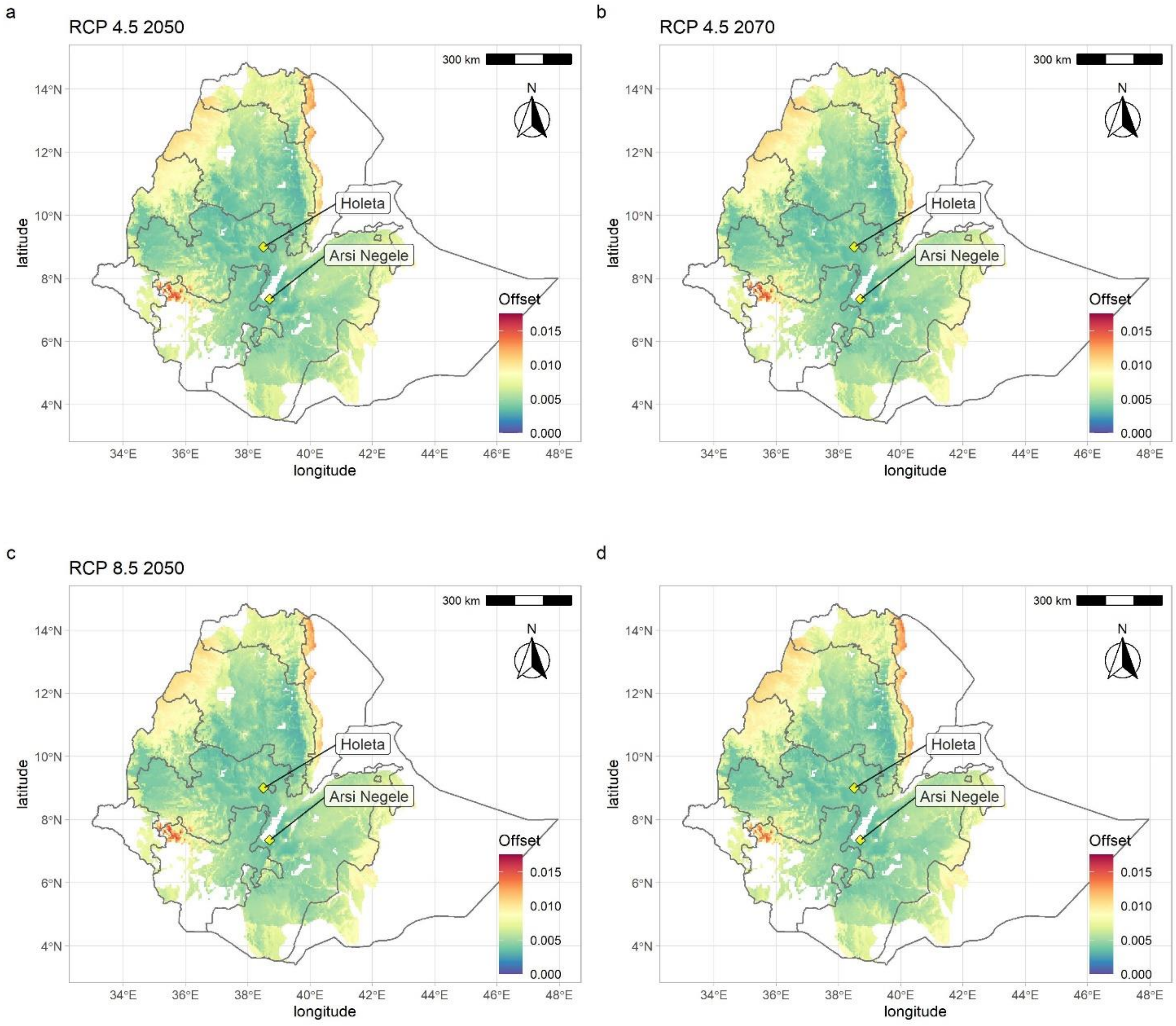


**Figure S9.** Genomic vulnerability of barley across the cropping area based on ensemble modelling projections for two representative concentration pathways: RCP4.5 and RCP8.5 at the horizons of 2050 and 2070. The color scale indicates the magnitude of the mismatch (*i.e.* Euclidean distance) between current and projected climate-driven turnover in allele frequencies.
